# Towards an Integrated Map of Genetic Interactions in Cancer Cells

**DOI:** 10.1101/120964

**Authors:** Benedikt Rauscher, Florian Heigwer, Luisa Henkel, Thomas Hielscher, Oksana Voloshanenko, Michael Boutros

**Affiliations:** Division of Signaling and Functional Genomics, German Cancer Research Center (DKFZ) and Heidelberg University, 69120 Heidelberg, Germany; Division of Biostatistics, German Cancer Research Center (DKFZ), 69120 Heidelberg, Germany

**Keywords:** genetic interactions, networks, epistasis, synthetic lethality, cancer

## Abstract

Cancer genomes often harbor hundreds of molecular aberrations. Such genetic variants can be drivers or passengers of tumorigenesis and, as a side effect, create new vulnerabilities for potential therapeutic exploitation. To systematically identify genotype-dependent vulnerabilities and synthetic lethal interactions, forward genetic screens in different genetic backgrounds have been conducted. We devised MINGLE, a computational framework that integrates CRISPR/Cas9 screens originating from many different libraries and laboratories to build genetic interaction maps. It builds on analytical approaches that were established for genetic network discovery in model organisms. We applied this method to integrate and analyze data from 85 CRISPR/Cas9 screens in human cancer cell lines combining functional data with information on genetic variants to explore the relationships of more than 2.1 million gene-background relationships. In addition to known dependencies, our analysis identified new genotype-specific vulnerabilities of cancer cells. Experimental validation of predicted vulnerabilities associated with aberrant Wnt/β-catenin signaling identified *GANAB* and *PRKCSH* as new positive regulators of Wnt/β-catenin signaling. By clustering genes with similar genetic interaction profiles, we drew the largest genetic network in cancer cells to date. Our scalable approach highlights how diverse genetic screens can be integrated to systematically build informative maps of genetic interactions in cancer, which can grow dynamically as more data is included.

## INTRODUCTION

Genes rarely function in isolation to affect phenotypes at the cellular or organismal level. Many studies have described how genes act in complex networks to maintain homeostasis by fine-tuning cellular or organismal reactions to internal or external stimuli (Bergman & Siegal 2003). A loss of genetic buffering can result in the emergence of diseases such as cancer (Hartman et al. 2001; Hartwell et al. 1997). In turn, mutations can create genetic vulnerabilities in cancer cells, for example, by deactivating one of two genetically buffered pathways (Luo et al. 2009; Nagel et al. 2016; Torti & Trusolino 2011). Therapeutic approaches attempt to exploit such events by selectively inducing cell death in cancer cells while causing little harm to normal cells (Kaelin 2005; Nijman 2011).

To systematically identify genetic interactions, pairwise gene knockout or knockdown experiments can be performed (Mani et al. 2008). In cases where a measured fitness defect of the double mutant is stronger than expected based on the two single mutant phenotypes, the interaction is called aggravating or synthetic lethal (Bridges 1922). In contrast, a buffering (or alleviating) interaction is observed when the double mutant’s measured phenotype is weaker than expected. Arrayed screens, performed by mating of loss-of-function mutant yeast strains have pioneered combinatorial screening (Baryshnikova et al. 2010; Costanzo et al. 2010; Davierwala et al. 2005; Tong et al. 2001; Costanzo et al. 2016). Methods of pairwise gene perturbation were later extended using combinatorial RNA interference (RNAi) to map genetic interactions in cultured metazoan cells (Horn et al. 2011; Laufer et al. 2013; Fischer et al. 2015; Byrne et al. 2007; Snijder et al. 2013; Srivas et al. 2016). However, screening of all pairwise gene combinations scales poorly with increasing genome size and novel approaches are necessary to facilitate the generation of large genetic interaction maps of complex organisms while minimizing cost and experimental effort.

Genome-scale perturbation screens can now be efficiently performed in many cell lines using CRISPR/Cas9 (Barrangou 2014; Doudna & Charpentier 2014; Horlbeck et al. 2016; Shalem et al. 2015; Wang et al. 2014) or RNAi (Brummelkamp et al. 2002; Kampmann et al. 2013; Sims et al. 2011) for the targeted perturbation of genes by knockout or knockdown. Since each cell line has a different genetic background, this enables the investigation of genotype-specific vulnerabilities (Hart et al. 2015; Steinhart et al. 2017; Wang et al. 2017; Tzelepis et al. 2016; Garnett et al. 2012; Iorio et al. 2016; Martin et al. 2017; Tsherniak et al. 2017; McDonald et al. 2017). To describe a genetic interaction previous studies have mostly relied on the definition of ‘statistical epistasis’ introduced by R. A. Fisher (Fisher 1930). Here, a genetic interaction is defined as a statistical deviation from the additive combination of two loci in how they affect a phenotype of interest (Phillips 2008). This definition does not necessarily assume a standardized genetic background and thus provides a theoretical framework applicable to map genetic interactions in cancer cell lines despite the presence of additional confounding mutations. To leverage the community’s collective effort to functionally characterize cancer cell lines it is desirable to combine and analyze genetic screens of different origin in an integrated manner. This, however, is not easily put into practice as various sources of technical variation such as different sgRNA libraries or experimental protocols can affect the data and confound comparative analyses.

Here we propose a computational framework that integrates CRISPR/Cas9 screens of diverse origin to map genetic interactions in cancer cells. We apply this approach, which we termed MINGLE, to a curated data set consisting of 85 genome-scale CRISPR/Cas9 screens in 60 different human cancer cell lines generated in various different laboratories (Figure 1A). We first show that a two-step normalization approach can be applied to enable quantitative comparison of phenotypes derived from different screens (Figure S1A). We then demonstrate how concepts that have previously been applied to map genetic networks in model organisms can be adapted and applied to this dataset to score gene-gene combinations for genetic interactions. Combining the intrinsic profile of genetic alterations of each cell line present in the dataset with gene-level viability phenotypes we tested 2.1 million pairwise gene combinations by comparing wild-type against altered alleles in cell lines (Figure 1B-C). Using these predictions, we were able to identify new regulators of the Wnt/β-catenin signaling pathway. Our results suggest that the genes *PRKCSH* and *GANAB*, which together form the Glucosidase II complex, regulate the secretion of active Wnt ligands. Finally, we functionally clustered genes by the similarity of their interaction profiles and demonstrate that these profiles are informative predictors of functional gene similarity (Figure 1D). We generated a map of genetic interactions in cancer cells by connecting genes with similar profiles and identified network modules with similar functional characteristics.

**Figure 1.**
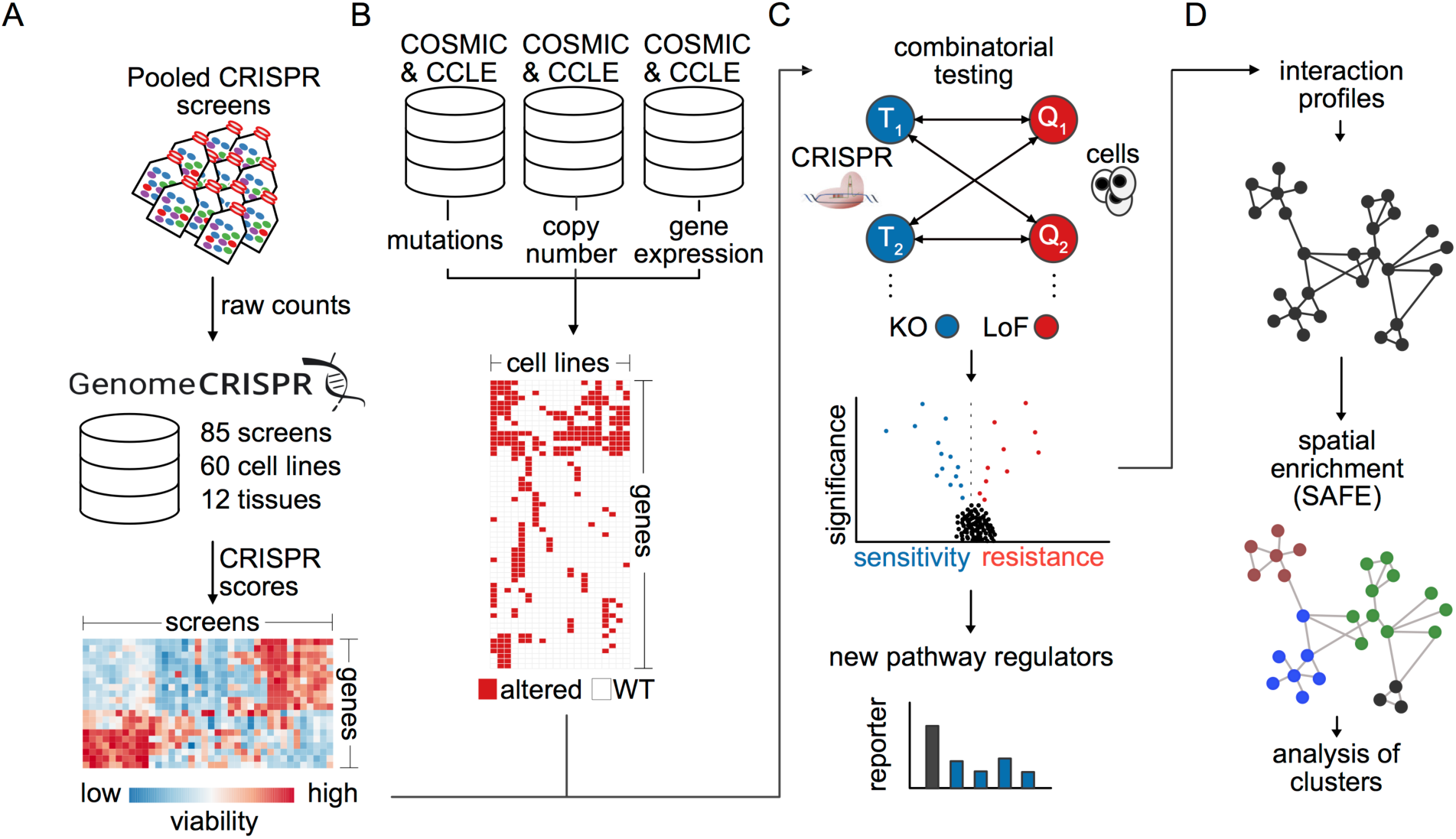
An integrated analysis approach to identify genetic interactions in cancer cells. (A) Data from CRISPR-Cas9 screens in 60 cancer cell lines were re-analyzed and integrated. The results were integrated into a global perturbation response profile. (B) Mutation, copy number and mRNA expression data from the COSMIC and CCLE databases were combined to create a map of genetic alterations across these cell lines. (C) To identify genetic dependencies between gene combinations that could shed light on the genetic wiring of cancer cells, perturbation response of more than 2.1 million gene-gene-combinations was examined to infer genetic interactions. (D) Interaction profiles were calculated for gene combinations based on the correlation of their interactions as determined by interaction scores (π scores). Spatial enrichment analysis was performed to identify functional modules in the network.

## RESULTS

### Integrating CRISPR/Cas9 phenotypes from different studies

In order to systematically predict interactions between genes knocked out by CRISPR/Cas9 and genes functionally impaired by mutations in cancer cells, we reanalyzed a set of 85 CRISPR/Cas9 viability screens in 60 cell lines (Figure 1A, Supplementary Table 3). These screens were performed in different laboratories and vary in terms of library and vector design as well as screening protocols. In order to integrate these data (Figure S1A), we first calculated gene level CRISPR scores (average log_2_ fold change of sgRNA abundance; Wang et al. 2017) individually for each screen. We then quantile-normalized the data to correct for systematic biases between screens as for example varying selection times can lead to differences in phenotypic strength. Examination of the resulting data set revealed considerable batch effects driven primarily by the sgRNA library used for screening (Figure S1B). These batch effects appeared to be non-systematic differing from gene to gene. For example, cyclin-dependent kinase 7 (*CDK7*) is a gene known to play important roles in both, cell cycle progression and transcription (Fisher 2005), and is expected to be a broadly essential gene (Hart et al. 2017). Accordingly, knockout of *CDK7* consistently led to decreased viability in the majority of experiments. The screens in which no viability phenotype was observed upon *CDK7* knockout were all conducted using the same library (Figure S1C). Since the cell lines screened with this library are derived from various different tissues and cancer types and a common resistance to *CDK7* knockout seems unlikely. A more probable explanation for the observed batch effect might be the inability of *CDK7* targeting sgRNAs in this library to generate a knockout in the first place. If not considered and corrected, such batch effects can introduce false predictions (Figure S1D), underlining the requirement of an efficient strategy for their adjustment. To this end, we hypothesized that a gene knockout should, on average, have the same effect across screens, regardless of the library used. We then applied a model-based approach to systematically scan for potential batch effects where the phenotypes generated by one library differed significantly (FDR < 5%) from the observed median phenotype across all libraries. In order to protect real biological effects, we used a robust linear model for testing, which is robust towards strong biological effects present in the data in the form of outliers. In cases, in which a significant difference between the phenotypes generated by one library and the median phenotype across all libraries could be detected, we performed an adjustment by subtracting the estimated difference between the library affected by the batch effect and the remaining libraries (Figure S1B). It is important to point out, that this approach can be inappropriate when there is a correlation between an sgRNA library and a biological covariate, for example if most cell lines screened with this specific library are derived from similar tissues. This is not the case for most libraries included in this analysis. For example, the GeCKOv2 and TKOv1 libraries have been used to screen a wide variety of cell lines derived from different tissues and cancer types (Aguirre et al. 2016; Hart et al. 2015; Steinhart et al. 2017). An exception, however, are the screens performed by Wang et al. (Wang et al. 2017) as well as Tzelepis et al. (Tzelepis et al. 2016). In these studies, screens were performed primarily in acute myeloid leukemia (AML) cell lines. In order to preserve such tissue-specific phenotypes through batch correction, our model-based approach allows to include biological covariates such as a cell line’s tissue or cancer type into the batch modelling, which can then distinguish between technical and biological variability. In order to validate our data integration approach, we performed a variety of quality control analyses. First, we clustered all screens based on the normalized CRISPR scores (Figure 2A, Figure S1F). In many cases, screens that were performed in different laboratories with different libraries but using the same cell line clustered together. Moreover, we observed a tendency for cell lines sharing the same tissue origin to group together. For example, we could identify distinct clusters of AML cell lines and adenocarcinoma cell lines. These results suggest appropriate correction of technical bias, leaving the biological variability across cell lines as the main driver of the clustering. We next assessed whether normalized CRISPR scores can be compared quantitatively across screens. Here, we randomly selected nine core-essential polymerases and plotted normalized CRISPR scores for these genes across screens (Figure 2B). CRISPR scores for essential polymerases were negative and approximately on the same level with no noticeable differences between screens published in different studies, suggesting that quantitative comparison of scores is indeed feasible and that expected negative viability phenotypes of core-essential gene knockouts are preserved throughout normalization. We wondered, if the normalization procedure could potentially introduce false phenotypes. Generally, this can be ruled out with the help of non-targeting controls, which, however, were not available for all experiments in our dataset. As a replacement, we therefore selected all screens performed in female cell lines and plotted normalized CRISPR scores for nine randomly selected genes located on the Y chromosome (Figure 2C). We observed CRISPR scores to be approximately 0, implying that no false phenotypes are introduced artificially by the normalization. Next, we determined how well core-essential and non-essential reference genes (Hart et al. 2015; Hart et al. 2017) could be separated based on the normalized CRISPR scores by generating precision-recall-curves (Figure 2D), based on which we observed good performance across all screens. We further examined if the normalized CRISPR scores could capture well-studied examples of oncogene addiction. Oncogene addiction describes a phenomenon where cancer cells, albeit harboring many molecular aberrations, become strongly dependent on only a single one of them. Reversing this abnormality leads to growth inhibition and apoptosis (Weinstein & Joe 2006). We selected the well-studied oncogenes *KRAS*, *NRAS*, *BRAF* and *PIK3CA* and compared the CRISPR scores of cell lines harboring a mutation of these genes to the rest of the cell lines (Figure 2E-H). As expected, we observed considerably stronger phenotypes in the mutated cells as compared to the wild-type cells. Last, we determined if genetic dependencies previously identified in screens used for our analysis could be reproduced (Figure S1E). In all cases, we could achieve comparable results to those previously published, corroborating the usage of normalized CRISPR scores for valid inter-screen-analysis.

**Figure 2.**
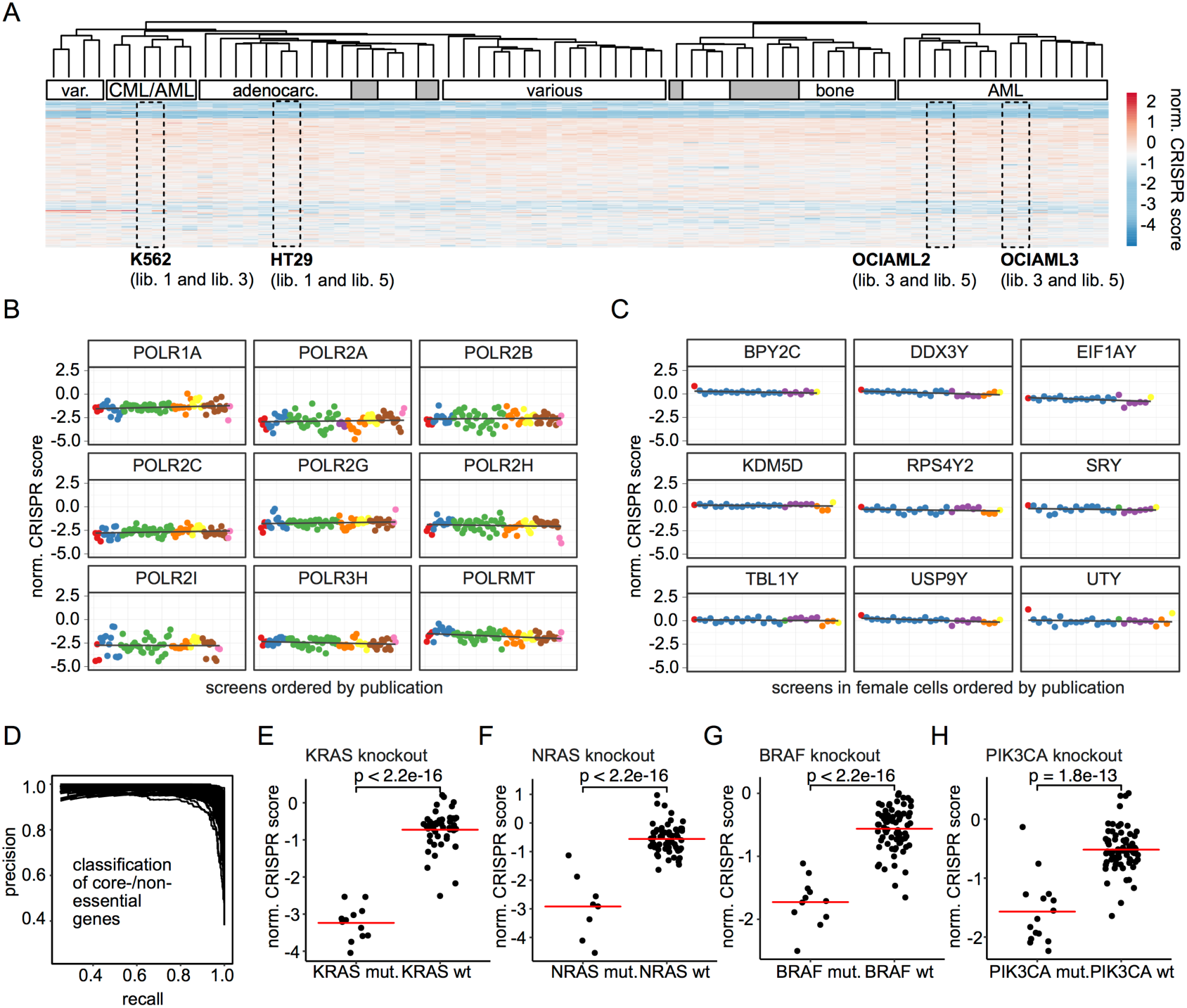
Results and quality control of data integration and normalization. (A) A heat map shows a clustering of normalized CRISPR scores (average log2 fold change of sgRNAs targeting a gene) for genes present in each sgRNA library used in screens included in the analysis. Rectangular windows highlight experiments where screens performed in the same cell line but in different laboratories cluster together. White annotation bars indicate shared biological properties of the cell lines in each cluster. Grey bars indicate the annotated cell line does not fit to the annotation of other cell line in the same cluster. (B) Normalized CRISPR scores across experiments are displayed for a randomly selected set of 9 core-essential polymerases. Each dot corresponds to one screen and different colors highlight the publications that the data were derived from. More negative CRISPR scores indicate a more negative viability response upon gene knockout. (C) Normalized CRISPR scores across experiments in female cell lines are displayed for a randomly selected set of 9 genes located on the Y chromosome serving as non-targeting controls. Colors depict different publications. (D) Precision-recall-curves showing the performance of normalized CRISPR scores at distinguishing core-essential from non-essential genes. Each line corresponds to one experiment. High recall while maintaining high precision indicates good performance. (E-H) Comparison of normalized CRISPR scores in a different genetic background for four different control dependencies.

### Interactions between gene knockouts and cancer alterations reveal genetic wiring maps

In order to determine genetic interactions, we formed all pairwise combinations between genes knocked out by CRISPR/Cas9 in pooled viability screens (target genes) and genes altered in cancer cells (query genes) (Figure 1C). We only considered genes as queries if they contain an alteration in at least three distinct cell lines (Supplementary Table 1). A cancer alteration was defined as a somatic mutation, a somatic copy number alteration (SCNA) or differential expression of a gene. We pooled alterations for each gene based on three assumptions: We assumed that (1) a loss of gene copy number behaves similarly to a disruptive somatic mutation (e.g. a frame shift mutation or a nonsense mutation), (2) a gain of copy number behaves similarly to a gain of gene expression and that (3) somatic mutations of the same gene have, on average, a similar functional consequence. Even though these assumptions, especially number 3, do in reality not always hold true, we found them to be a useful approximation judging by the results we obtained in downstream genetic interaction analyses. In addition, we further refined pooled alterations by manual curation excluding cell lines with alterations known to be functionally dissimilar to other alterations of the same gene. This, however, was only possible for well-characterized genes. In total, we formed 3.8 million gene pairs of 17,218 target genes and 221 query genes.

Assuming that two genes do in most cases not interact with each other, we first performed a statistical test for each gene pair, comparing normalized CRISPR scores of cells that contain an alteration of the query gene to cells that do not contain the alteration. Here, we used a multilevel model including the cell line corresponding to each data point as a random effect to account for biases that could potentially be introduced when one cell line was screened multiple times. In some cases, we observed high correlation between several query genes (Figure S2A). This observation can, for example, be explained by a co-deletion of genes that are located close to each other on the genome. For instance, *CDKN2A*, a tumor suppressor gene (Liggett & Sidransky 1998) located on chromosome band 9p21 is often co-deleted with its surrounding genes (Muller et al. 2015). In such cases, it is not possible to determine with which of the two potential query genes a target gene should be predicted to interact. We addressed this by aggregating identical query genes, as determined by the correlation of their model coefficients, into ‘meta genes’ that we then used for downstream analyses (Figure S2B). To quantify the interaction strength of each gene pair, we calculated π-scores (Figure 3A-B) as described previously (Horn et al. 2011; Laufer et al. 2013; Fischer et al. 2015). Altogether, our analysis predicted 17,545 gene-gene-interactions at FDR < 0.2 (0.8% of total combinations tested after meta gene aggregation).

**Figure 3.**
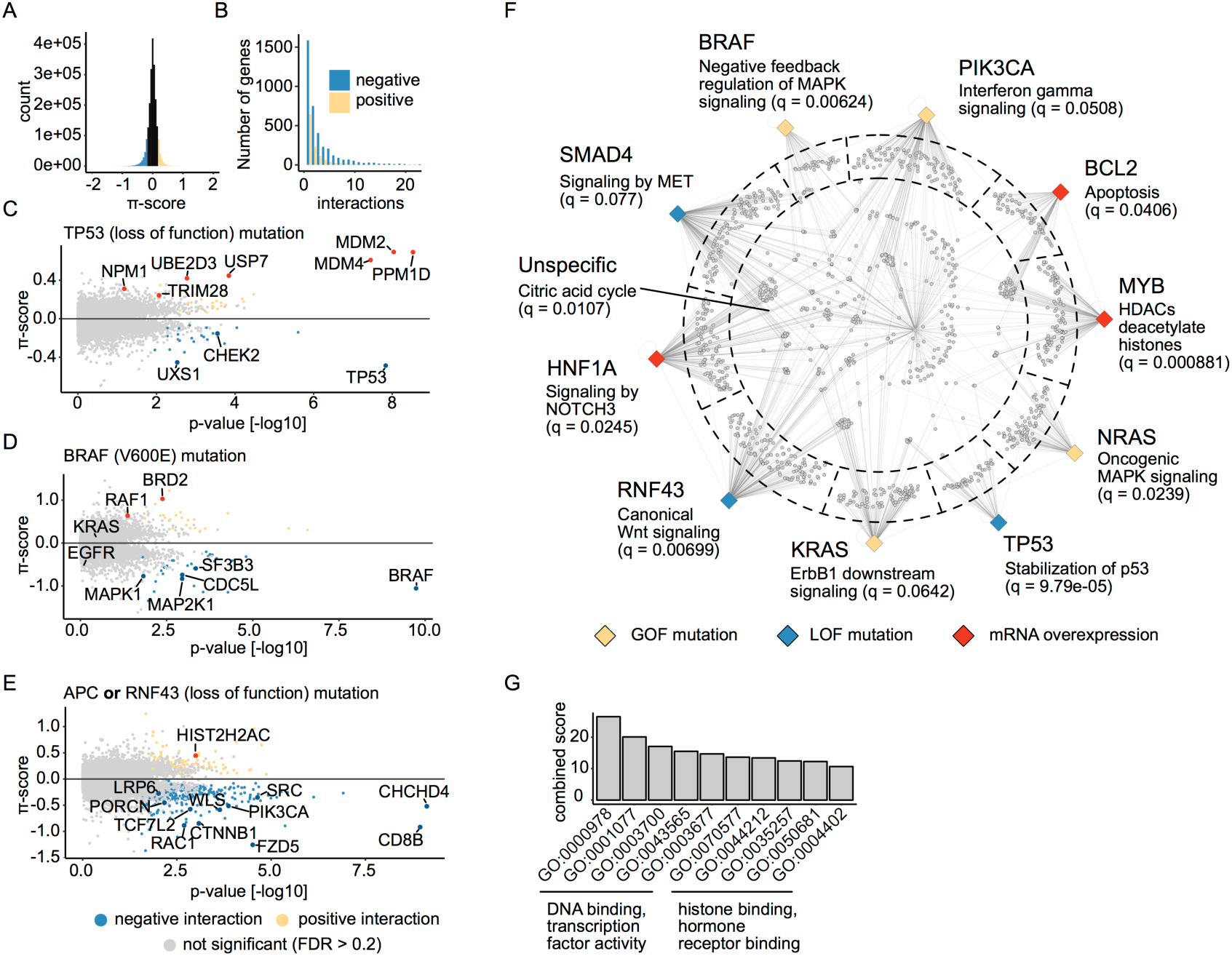
Results of predicted genetic interactions. (A) Distribution of π-scores calculated for each pairwise interaction. Negative values indicate negative (aggravating interactions) and positive values indicate positive (buffering) interactions. Values greater than 0.2 and less than −0.2 are colored yellow and blue, respectively. (B) The number of positive and negative interactions per gene. Interactions with a π-score greater than 0.2 are considered positive and interactions with a π-score of less than −0.2 are considered negative. (C-E) Volcano plots showing genes interacting with TP53 loss-of-function mutations (C), BRAF V600E mutations (D) and APC or RNF43 loss-of-function mutations (E). Each dot corresponds to one gene. Interactions that are significant at FDR < 0.2 are colored in blue in case the interaction is negative or yellow if it is positive. Selected genes are highlighted and labeled. (F) A network graph showing gene set enrichment results for sets of interaction partners. Each of the colored diamonds corresponds to one of 10 selected query alterations. The color of each diamond indicates the type of alteration as described in the legend at the bottom. Each grey dot connected to one or more query gene nodes represents a target gene that interacts (FDR < 0.2) with the query. Gene set enrichment analysis was performed for genes that fall in the same compartment as indicated by the dashed line. Genes in compartments towards the edge interact with one specific query. Genes positioned in the center of the circle have a more promiscuous interaction profile. Selected enriched pathway terms are used to label the query gene nodes. (G) GO terms enriched among 40 query genes with the most interactions (|π| > 0.2, FDR < 0.2).

Examining the proposed interactions, we found that our analysis was able to recover many previously characterized dependencies across several pathways that have been extensively studied in the past (Figure 3, Figure S2F-H). For example, we identified many positive interactions (i.e. cells containing an alteration of the query gene are more resistant to perturbation of the target gene) between *TP53* and several genes involved in stabilization of the p53 protein (Figure 3C). In wild-type cells, p53 is kept at low abundance by E3/E4 ubiquitin ligases including for example *MDM2* and *MDM4* (Figure S2G), which can mediate its degradation via the proteasome (Lavin & Gueven 2006; Frum & Grossman 2014). Knockout of these ubiquitin ligases likely leads to an accumulation of p53, which might then mediate apoptosis and impede proliferation resulting in a negative viability phenotype. In tumor cells, missense mutations of the *TP53* gene can inhibit p53 degradation (Frum & Grossman 2014; Lavin & Gueven 2006) where it can accumulate and act as an oncogene (Oren & Rotter 2010), which could explain the resistance of *TP53* mutated cell lines to E3/E4 ubiquitin ligases. An interaction that at first glance might seem surprising is a negative interaction of *TP53* with itself (i.e. cells with a *TP53* mutation are more sensitive to *TP53* knockout). In the context of epistasis, however, this might be explained by the fact that in *TP53* wild-type cells, where *TP53* acts as a tumor suppressor, its knockout leads to a gain of viability phenotype, which is not the case for tumor cells which already harbor mutations in *TP53* (Figure S2H). Next, we looked at predicted interactions of the *BRAF* oncogene. Unsurprisingly, we found negative interactions with *BRAF* itself as well as *MAP2K1* (MEK1) and *MAPK1* (ERK2), both of which lie downstream of *BRAF* in the MAPK signaling cascade (Seger & Krebs 1995). In contrast, no interactions were found for upstream components of the pathway such as *KRAS* or *EGFR* (Figure 3D), likely because the constitutive activation of *BRAF* caused by its mutation confers independence on upstream pathway components. Following previous studies (Brockmann et al. 2017), we reasoned that genes that interact specifically with one or few related query genes should be functionally related. We thus selected ten query genes including their predicted interaction partners at FDR < 20% and performed gene set over-representation analysis (Kamburov et al. 2013) for groups of target genes specifically interacting with one of the selected queries (Figure 3F). Looking at pathways over-represented within the analyzed set of genes, we found several well-characterized relationships linking for example mutations of *KRAS*, *NRAS* or *BRAF* to MAPK signaling, *BCL2* to apoptosis or *TP53* to the stabilization thereof, suggesting a high number of true predictions. In addition, our analysis proposes genetic interactions for many other less well-studied query genes (a full list of predicted interactions can be found in Supplementary Table 4). To find traits shared between query genes for which high interaction numbers were predicted (Figure S2E), we performed GO (Ashburner et al. 2000) molecular function enrichment analysis (Kuleshov et al. 2016). Unsurprisingly, we found that GO terms with the highest enrichment scores were related to transcription factor activity (Figure 3G). Other high-ranking GO terms were related to chromatin remodeling and hormone receptor binding.

We hypothesized that it should be possible to combine functionally related query genes in order to improve prediction of regulators of signaling pathways. Consequently, we combined loss of function mutations of the genes *APC* and *RNF43* (Supplemental Table 2) into a ‘Wnt mutation’ query meta-gene. Both, *APC* and *RNF43*, are potent and frequently mutated negative regulators of the Wnt/β-catenin signaling pathway (Tsukiyama et al. 2015; de Lau et al. 2014; Zhan et al. 2017; Polakis 2012) - a pathway that is aberrantly regulated in various cancers (Zhan et al. 2017; Polakis 2012; Giannakis et al. 2014). In the absence of Wnt ligands, APC regulates β-catenin activity *via* the formation of a destruction complex with GSK3β and Axin1, which mediates β-catenin phosphorylation. Phosphorylated β-catenin is targeted for degradation by the proteasome. Binding of canonical Wnts to Frizzled receptors and LRP5/6 co-receptors on the cell surface inhibits the formation of the destruction complex, which results in β-catenin stabilization and its translocation to the nucleus. Within the nucleus, β-catenin interacts with TCF/LEF transcription factors and activates transcription of Wnt target genes, which mediate cell growth and survival (MacDonald et al. 2009). *RNF43* is an E3 ubiquitin ligase that can induce ubiquitination and subsequent degradation of the Wnt-Frizzled complex (MacDonald et al. 2009; Clevers & Nusse 2012), thus inhibiting β-catenin signaling. Consequently, disruptive mutations in *APC* or *RNF43* can promote activation of the pathway. Examining genes predicted to interact with loss-of-function mutations of either *APC* or *RNF43*, we observed many known regulators of Wnt/β-catenin signaling (Figure 3E). Among these we identified for example regulators of Wnt ligand secretion, *TCF7L2* and *CTNNB1* which together form the TCF/β-catenin transcription factor complex, and other genes, which have previously been linked to the Wnt/β-catenin pathway (Chen et al. 2014; Ormanns et al. 2014).

### Dependency analysis of Wnt pathway alterations reveals novel regulators of Wnt/β-catenin signaling

We hypothesized that among known modulators of Wnt/β-catenin signaling, our analysis should also identify so far uncharacterized pathway regulators. Inactivating mutations of the *RNF43* gene, for example, have previously been shown to confer dependency on Wnt/β-catenin signaling (Jiang et al. 2013; Steinhart et al. 2017) so we reasoned that negative interactions of *RNF43* could point to positive pathway regulators. Besides known Wnt pathway regulators our analysis revealed negative interactions between RNF43 and several interesting target genes (Supplementary Table 4). We aimed to experimentally validate these predictions and proceeded by selecting three high-scoring candidate genes reported to be involved in protein glycosylation (D’Alessio & Dahms 2015) for follow-up (Figure 4A). Two of these genes, *PRKCSH* and *GANAB*, together form the heterodimeric Glucosidase II. The third candidate, *UGP2*, is involved in carbohydrate synthesis (Wang et al. 2016). We knocked down each of the candidate genes using at least three different siRNAs (Figure 4B, Figure S3B, Materials & Methods) or a pool consisting of the same reagents in HEK293T cells (Figure 4B). HEK293T cells were chosen as a non-tumorigenic, well-established model for canonical Wnt signaling activation, which harbor no known mutations in the Wnt pathway. Furthermore, HEK293T feature an inactive state of canonical Wnt signaling, which is why the pathway can be activated by overexpression of some of its key components (Wnt3, Dvl3 and β-catenin). Overexpression of Wnt3 mimics auto-paracrine activation of canonical Wnt signaling at the level of the Wnt secreting cell, while overexpression of Dvl3 induces the pathway downstream of the receptor complex in the receiving cells. Overexpression of β-catenin leads to its overload and thus stabilization in the receiving cells and activation of the pathway downstream of APC (Figure 4B, Figure S3A). We observed, that knockdown of each of the tested candidate genes followed by pathway activation induced by Wnt3 expression resulted in strongly reduced activation of a TCF4/Wnt reporter, which mimics transcription activation of genes regulated by β-catenin (Figure 4B). Interestingly, knockdown of *GANAB*, *PRKCSH* or *UGP2* did not show a strong effect on reporter activity or even enhanced induction upon transfection with Dvl3 or β-catenin expression plasmids (Figure 4B). These results allow to conclude an interference of the candidates investigated at the level of Wnt secretion or at the receptor level, since the negative effect on Wnt activity is abolished upon further downstream pathway activation by Dvl3 or β-catenin. To further investigate the role of the Glucosidase II complex and by this protein glycosylation, secretion and quality control of glycoprotein folding in the ER in the context of Wnt signaling, we performed a Wnt secretion assay upon knockdown of *PRKCSH* and *GANAB* (Figure 4D; D’Alessio & Dahms 2015). For this we coupled Wnt3 to a NanoLuciferase (Hall et al. 2012) sequence within a Wnt3 expression plasmid. The NanoLuciferase sequence was integrated either after the signal peptide (NLucWnt3) or at the C-terminus of Wnt3 (Wnt3NLuc) to exclude an effect of NanoLuciferase coupling on Wnt3 secretion. A NanoLuciferase readout subsequently allowed to detect secreted Wnt3 proteins in the cell culture supernatant and to normalize it to the amount of Wnt3 in the cell lysate. Upon knockdown of either GANAB or PRKCSH, Wnt3 secretion was reduced about 40-50% using either the NLucWnt3 or Wnt3NLuc constructs (Figure 4C, Figure S3C). These data substantiate an already published necessity of Wnt ligand glycosylation for successful secretion of Wnt proteins (Figure 4D; Komekado et al. 2007).

**Figure 4.**
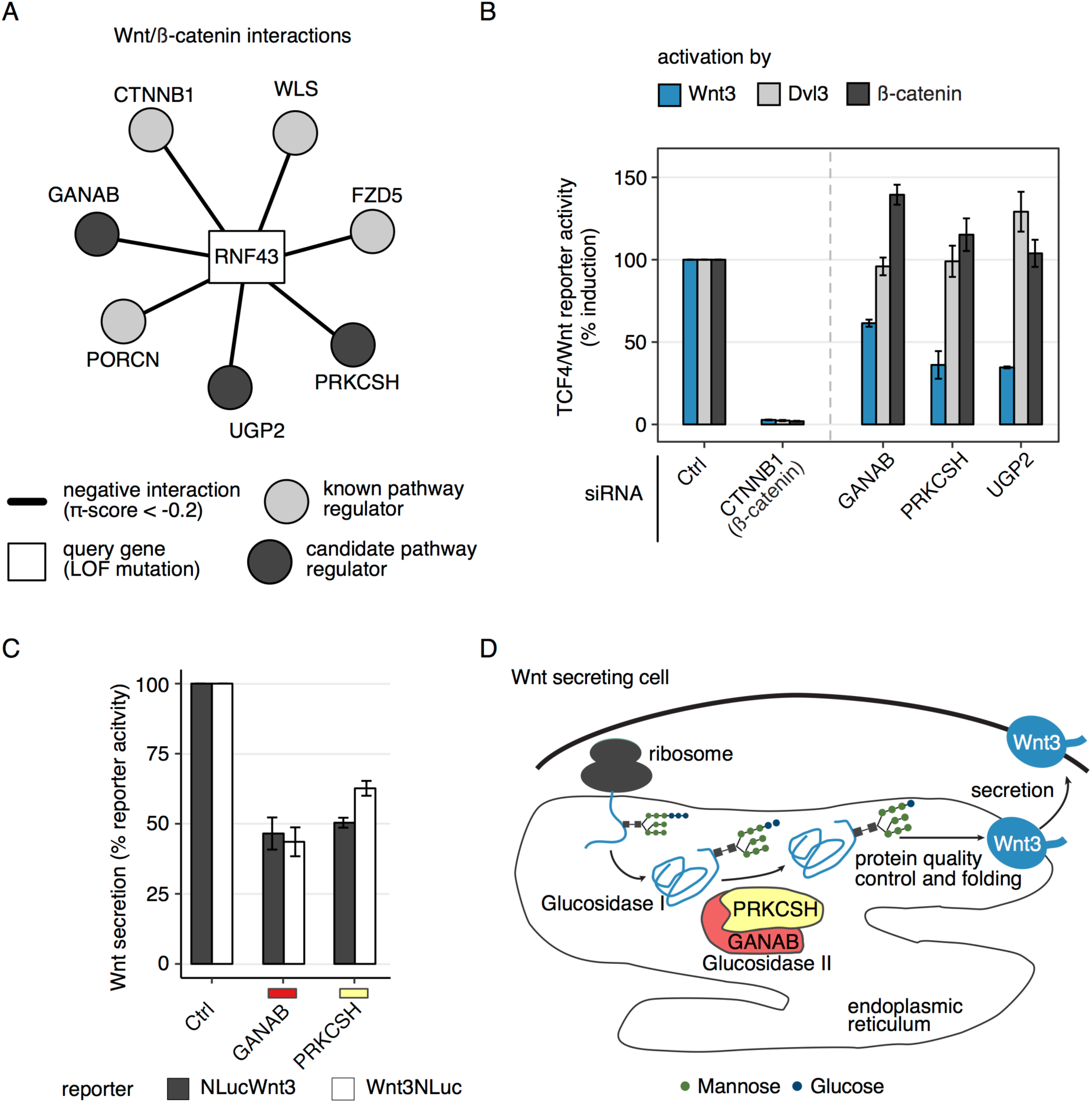
Candidate genes *GANAB* and *PRKCSH* regulate Wnt secretion. (A) Three candidate genes (dark gray circles) interact with the *RNF43* query gene (rectangle), similar to well characterized pathway components (light grey circles). (B) HEK293T cells were reverse transfected with siRNA pools targeting genes labeled on the x-axis. 24 hours after transfection Wnt signaling was activated by overexpression of Wnt3, Dvl3 or β-catenin plasmids. The TCF4/Wnt Firefly luciferase signal was normalized to the actin-*Renilla* signal. Results are shown as averages of 3-4 independent experiments ± s.e.m. (C) HEK293T cells were reverse transfected with pooled siRNAs targeting *GANAB* or *PRKCSH*. After 24 hours the indicated Wnt3 NanoLuciferase constructs were transfected together with a CMV Firefly luciferase reporter. 48 hours later Luciferase signals were measured in the medium and lysate. % reporter activity denotes the Wnt3 NanoLuciferase signal in the medium normalized to NanoLuciferase and Firefly luciferase signals in the lysate. Results are shown as averages of 3 independent experiments ± s.e.m. (D) Schematic depiction of a hypothetical mechanism where Wnt3 secretion is controlled by Glucosidase II.

### Similarity of interaction profiles predicts functional relationship of genes

Several studies have previously shown that functionally similar genes can be identified by comparing their interaction profiles. Here, the vectors of interaction scores across query genes are compared for all possible pairs of target genes using a measure of similarity - most commonly their correlation. Two target genes with highly correlating interaction profiles are then predicted to share biological function through guilt by association (Figure 1D). Encouraged by the observation of pathway enrichment among target genes predicted to interact with the same query we reasoned that an analysis of interaction profile similarity should also be possible based on our results despite a relatively low number of query genes (167 after aggregation of highly similar query genes). Consequently, we correlated Pearson correlation coefficients of π-score interaction profiles for all pairwise combination of target genes. We reasoned that data about known protein complex co-membership should be able to serve as a reference to estimate the predictive power of our approach. Hence, we downloaded all human protein complex data from the CORUM (Ruepp et al. 2010) database and compared our predicted associations to the known protein complex data by receiver operator characteristic (ROC) analysis. Initially, this analysis revealed our predictions of protein complex co-membership to be unsatisfactory. After careful inspection of the predicted relationships we noticed that the correlation coefficient was in most cases considerable influenced by very small π-scores. Such data points do not hold a lot of biological information as they merely indicate that there might be no connection between a target and a query gene based on a viability phenotype. Hence, we hypothesized that excluding interactions with very low π-scores should shift more weight onto more informative data points and should therefore lead to more meaningful predictions of co-functionality. We consequently excluded all interactions with π-score < 0.2 and repeated the above analysis. As excluding interactions with a low π-score violates the Pearson correlations’ assumption of normality we used the non-parametric Spearman correlation instead. We calculated this correlation for all pairs of target genes where at least 5 pairwise complete data points were available. Repeating the ROC analysis as described above revealed a considerable improvement of the resulting predictions leading to results clearly better than random assignment (Figure 5A). In order to identify the most suitable parameter thresholds we systematically repeated this analysis using different combinations of the π_min_ (minimum π-score to be considered) and n_min_ (minimal number of pairwise complete data points) parameters. We noticed that more conservative parameter thresholds lead to higher performance at predicting protein complexes. However, the more conservative these thresholds become the more genes have to be excluded from the analysis due to insufficient data. Therefore, we decided to select π_min_=0.2 and n_min_=15 as parameters for downstream analyses, assuming these cutoffs to present a good compromise between the predictive power of the analysis and the number of genes that can be considered. Based on these parameters we found that our analysis holds power to correctly associate many closely interacting genes, such as for example *CTNNB1* and *TCF7L2* which together form the TCF/β-catenin transcription factor complex (Morin et al. 1997) or the *WNT10A FZD5* ligand receptor complex (Voloshanenko et al. 2017; Figure 5B). Similar interaction profiles could also be found for several members of the Mediator complex, a multi-subunit complex important for the transcriptional regulation of RNA polymerase II (Figure 5C).

**Figure 5.**
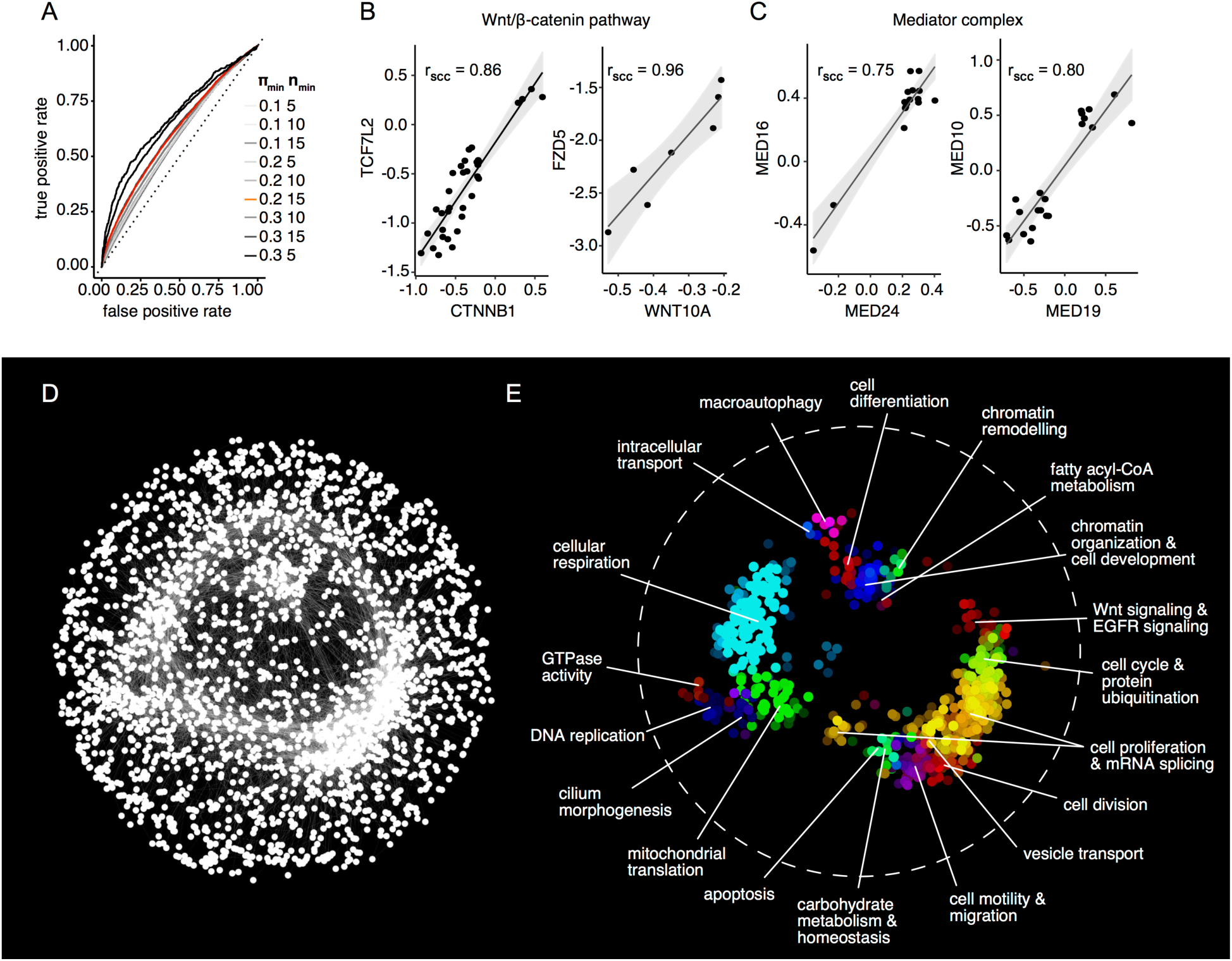
Highly correlating interaction profiles can predict functional similarity. (A) ROC curve displaying the performance of interaction profile similarity at predicting protein complex co-membership. Curves are shown for different filtering parameter combinations. The curve corresponding to the parameter combination used for downstream analysis (π_min_ = 0.2; n_min_ = 15) is highlighted in red. A grey dashed line indicates the performance expected by random assignment. (B-C) Examples of protein complexes where complex members display highly correlated interaction profiles (SCC = Spearman correlation coefficient). (D) Network of genes with highly correlated interaction profiles. 2,497 nodes (genes) are connected by 19,044 links (Bonferroni corrected p-value of the Spearman correlation of interaction profiles < 0.5). An edge-weighted spring embedded layout was used to position the nodes. (B) Spatial enrichment analysis with the SAFE algorithm highlights network modules consisting of genes with similar functional annotations based on Gene Ontology biological processes. The labels in the figure summarize the GO terms associated with each module.

We selected all target gene pairs for which the Bonferroni-corrected asymptotic p-value of the profile similarity (Spearman correlation) was smaller than 0.5 and connected them to a network, applying a force-directed spring-embedded layout that can position highly similar genes proximal to each other (Figure 5D). We next used Spatial Analysis of Functional Enrichment (SAFE; Baryshnikova 2016b; Baryshnikova 2016a) to identify regions in the network enriched for specific biological processes as annotated by Gene Ontology (GO; Ashburner et al. 2000; Figure 5E; Supplementary File 1). SAFE analysis revealed clustering of 19 sub-networks, which were associated with x different GO terms and comprised in total y genes.

In order to ensure that the observed modules do in fact resemble biologically meaningful functional clusters and are not just random artefacts of the analysis, we performed a random permutation analysis (Figure S4A-C). As expected, we observed that upon random reshuffling of links while keeping the genes and edge number the same the network loses its modular structure, resulting in one big cluster of genes in the center of the network. SAFE analysis reveals that this cluster enriches for metabolism genes, indicating that there is a general overrepresentation of metabolism genes among genes found to behave differentially in cancer cells.

Functionally enriched clusters not only cover biological processes commonly found to be implicated in cancer (e.g. “cell devision”, “Wnt & EGFR signaling” or “cell differentiation”) but also processes of general importance in cellular development and behavior (e.g. “cilium morphogenesis”, “intra cellular transport” and “macro autophagy”). This implicates that the approach presented here is indeed capable of identifying novel regulators of known pathway assemblies and previously uncharacterized members of know functional biological processes. This way we created an unprecedented resource of functional gene clusters to be exploited by future studies for deeper understanding of novel mechanisms influencing known bioprocesses, not only important in cancer but covering a wide range of biology. This resource can also be used to validate prior assumption of gene functions in any functional study. We anticipate that as data in more cell lines and phenotypes become available this functional map of a cell will continue to grow and improve.

## DISCUSSION

To identify novel functions of known genes or to assign cellular function to unknown genes, forward genetic screens have been conducted in many model systems ranging from bacteria to human cells (Boutros & Ahringer 2008). Combining high-throughput screening methods with the ability to reliably knock out every gene in the human genome by programmable nucleases now opens up the possibility of studying the consequences of complete or partial loss-of-function mutations with unprecedented accuracy in various mutational backgrounds. Genome-wide screens, predominantly for gene essentiality, have been performed and have identified a large number of known, new and context-specific essential genes (Evers et al. 2016; Hart et al. 2015; Morgens et al. 2016; Rauscher et al. 2017; Wang et al. 2014; Wang et al. 2015; Zhan & Boutros 2016). We developed a computational approach to integrate dozens of high-throughput CRISPR/Cas9 viability screens independent of screen size, library, Cas9 type and screening protocol. Because, compared to other techniques, CRISPR/Cas9 screens have shown to be a more sensitive method by which perturbation-induced phenotypes can be discovered in human cells (Hart et al. 2015; Wang et al. 2015), such an approach shows great promise for the systematic discovery of cancer vulnerabilities. We developed MINGLE, a computational framework that integrates CRISPR/Cas9 screens of diverse origin to map genetic interactions in cancer cells. We applied this approach to integrate data from 85 screens in human cancer cell lines and analyzed the viability effects of CRISPR/Cas9 perturbations in the context of the cell lines’ genetic backgrounds. By systematically evaluating 2.1 million combinations of genes, we uncovered genetic wiring maps including many known and novel dependencies between genes implicated in tumorigenesis and resistance to therapy. We further show that these maps can identify new regulators of pathways that play important roles in specific cancer types, e.g. β-catenin-dependent Wnt signaling.

Here we demonstrate that members of the Glucosidase II complex that our analysis identified as RNF43 interacting genes control signaling activity by regulation of Wnt3 ligand secretion, probably mediated by protein N-glycosylation. N-linked glycosylation is an ER-based process essential for protein secretion and folding (Xu & Ng 2015; Figure 4D). Whereas N-linked glycosylation of Wnt-3a has already been described in the past (Smolich et al. 1993), the importance of Wnt ligand glycosylation for secretion and pathway activation is controversially discussed. While some authors state a clear correlation between Wnt ligand glycosylation and secretion in a human cell line (Komekado et al. 2007), others could not observe loss of protein secretion upon suppressing protein N-glycosylation in *Drosophila* (Tang et al. 2012; Herr & Basler 2012). Our results support a role of three genes involved in protein glycosylation on Wnt pathway activation, which could be further supported by a reduction of Wnt ligand secretion upon knockdown of *GANAB* and *PRKCSH*.

Traditionally, genetic interactions have been examined by simultaneous perturbation of two genes. Our analysis is based on the idea that one of these perturbations can be mimicked by genetic alterations that naturally occur in cancer cells. Even though we find that this concept can indeed be applied to efficiently identify true interactions it poses a number of challenges. First of all, genetic alterations of each gene have to be pooled demanding certain assumptions about the similarity of their functional consequences. In nature, however, these assumptions do not always hold true which can confound the analysis. In this study we have attempted to address this issue by dividing alterations into logical groups, for example by pooling nonsense mutations and frameshift mutations as loss-of-function variants. We have further refined these annotations by manual curation excluding cell lines with variants known to be functionally distinct from others. Although this is currently only possible for well-characterized genes and we are confident that future advances regarding the functional characterization of cancer variants will greatly benefit our approach. Another challenge is posed by the fact that some genetic alterations are correlated because they co-occur in the same cell lines or cancer types. An example is the deletion of the chromosome 9p21 locus where the tumor suppressor *CDKN2A* is located. *CDKN2A* is often co-deleted with its neighboring genes (Muller et al. 2015) and it is thus not easily possible to understand which of them is the true driver behind a proposed interaction. This can further introduce a bias into the genetic similarity network. In our study we address this by aggregating fully correlated query genes into ‘meta genes‘ that we then proceed to use to calculate interactions and generate the genetic similarity network. To avoid bias we further calculate correlations of genetic interaction profiles based on only a subset of query genes such that no two query genes are more than 70% similar in terms of their cell line composition.

It has previously been demonstrated that profiles of synthetic genetic interactions can group functionally related genes through “guilt by association”. Studies in human cells have formerly relied on RNA interference. However, it has been shown that this method has limitations, such as off-targeting and dosage compensation effects, that can be overcome by CRISPR/Cas9. Our approaches allowed us to analyze interaction profiles using data from many high throughput CRISPR/Cas9 experiments. These profiles hold power to predict functional relationships of genes as we show by benchmarking against the CORUM protein complex database. As physical protein interactions as they occur in protein complexes represent only a subset of possible functional relationships we believe that this benchmarking can be interpreted as a lower bound for the predictive power of the analysis. We created a network that groups genes into clusters with enriched functional profiles. Findings from this analysis may be important for two reasons: first, hypotheses about the function of weakly characterized genes that are frequently deleted in cancer cells can be generated by looking at the common interaction partners members within functional network modules; and second, such a network may serve as a powerful tool to infer the function of entirely uncharacterized genes based on the function of connected genes. For example, over 10% of the genes in our network are not annotated with GO biological processes.

In its current state, a limiting factor of this type of analysis is the amount of available data. At present, there are approximately 200 genes that have been found to be frequently altered in the cell lines included in our data and for which synthetic genetic interactions can be tested. Therefore, only genes that interact with these genes can currently be examined. Nevertheless, this number will improve rapidly as new data are published, which will then allow for the creation of increasingly complex interaction networks. Pooling functionally related alterations of different genes as we demonstrate at the example of *RNF43* and *APC* can further expand the set of possible query genes. Our approach is scalable and can easily be expanded as new data becomes available. All in all, we believe that the presented approach can be a powerful way to systematically discover synthetic genetic interactions that may be of clinical interest. Furthermore, we believe that it can serve as an important asset to the quest towards more complete understanding of how human genes function. The presented workflow scales well as increasing amounts of data are becoming available.

We expect many more CRISPR/Cas9 screens in various cell lines to be carried out in future. We will expand our analysis once these data become available to improve and diversify our findings. Finally, we aim to extend our analysis to also include data from other experiment types such as physical interactions derived from protein-protein interaction studies. Most synthetic genetic interactions, for example, do not link genes that are members of the same pathways but instead they connect members of two interacting pathways (Kelley & Ideker 2005). Therefore, integrating synthetic interactions and physical interactions derived from protein-protein-interaction experiments might provide important new insights into how biological pathways interact with each other.

We further aim to make the predicted interactions available for browsing and download through the GenomCRISPR database, as we believe that they can be a useful resource to inform candidate gene selection for experiments that cannot be carried out at a genome-wide scale. These include, for example, *in vivo* screens in genetically engineered mouse models that are often limited by the number of cells that can be transfected or pairwise perturbation experiments as they are now conducted in human cells using CRISPR/Cas9 (Shen et al. 2017; Du et al. 2017), which are limited, by the number of possible gene combinations.

## METHODS

### Genetic profiles of cancer cell lines

To generate profiles of genetic alterations in GenomeCRISPR (Rauscher et al. 2017) cancer cell lines we relied on data publicly available in the COSMIC Cell Lines Project (Forbes et al. 2017), the Cancer Cell Line Encyclopedia (CCLE; Barretina et al. 2012) and additional data published previously by Bürckstümmer et al. for the KBM7 cell line (Bürckstümmer et al. 2013) and Klijn et al. (Klijn et al. 2014) (Figure 1B). Taken together, these data can characterize all except for 2 (a patient derived Glioblastoma cell lline and the RPE1 cell line) cell lines currently included in GenomeCRISPR. In total, 60 different cell lines were included in the analysis. For each of these cell lines a list of altered genes was generated, taking into consideration the following types of alterations: 1) gain of copy number events, 2) loss of copy number events, 3) somatic mutations, excluding silent mutations and in-frame insertions or deletions, and 4) mRNA overexpression.

### Selection of copy number alterations

First, copy number data was downloaded from the COSMIC Cell Lines Project v81, the CCLE (file dated 27-May-2017) and the Klijn et al. publication (Klijn et al. 2014). Gain and loss of copy number status was determined for each gene as follows: COSMIC provides a label for each copy number event that indicates whether the event can be classified as a gain or loss of copy number event. We adopted this classification for our analysis. In the paper by Klijn and colleagues, amplification and deletion of a gene was defined as > 1 or < −0.75 of the ploidy corrected copy number (Mermel et al. 2011; Klijn et al. 2014). Consequently, the same thresholds were used in our approach. Finally, CCLE provides log_2_-transformed copy number fold changes between healthy samples and cancer cell lines at the gene level. The absolute copy number of each gene per cell line was estimated from the fold change data as

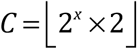

Where *C* is the absolute copy number and *x* is the log_2_ fold change between cell line and healthy sample. In order to assess whether this provides a realistic estimate of the total copy number we analyzed the derived copy number for all Y-chromosome genes in female cell lines where copy numbers of 0 were robustly estimated. Finally, we downloaded pre-processed gene-level copy number data from COSMIC. All genes where a copy number of 0 was estimated in a cell line were marked as loss-of-function genes. Copy number alteration events that were observed robustly across at least 2 different data sources were kept for downstream analysis after excluding alterations on the X and Y chromosomes.

### Selection of somatic mutations

Somatic mutation data were downloaded from COSMIC Cell Lines Project (version 81), the CCLE (Oncomap3 mutations dated 10-Apr-2012 and Hybrid Capture mutations dated 05-May-2015) and the Klijn et al. and Bürckstümmer publications. Missense mutations and frame-shift mutations were selected and mutations reported in disagreement between individual data sources were excluded. Next, missense mutations were classified into driver and passenger and driver as proposed by Anoosha et al. (Anoosha et al. 2016). Putative passenger mutations were excluded and the remaining mutations were kept for downstream analysis. After pooling copy number alterations and somatic mutations, we kept all genes as query genes where an alteration was observed in at least 3 different GenomeCRISPR cell lines.

### Selection of overexpressed genes

In order to define genes that are overexpressed in cell lines included in GenomeCRISPR, RMA (Irizarry et al. 2003) normalized microarray mRNA expression data were downloaded from CCLE (CCLE_Expression_2012-09-29.res dated 17-Oct-2012) and the COSMIC Cell Lines Project (v81). ComBat (Leek et al. 2012) was used to remove batch effects between the two different data sources and expression levels for cell lines featured in both sources were aggregated by computing the mean. Next, gene expression Z-scores were computed for each gene in each cell line. Genes on the COSMIC list of cancer census genes for which a Z-score > 2 was observed in at least 5 different GenomeCRISPR cell lines were kept for downstream analysis.

### Analysis of CRISPR-Cas9 screens

To compare viability phenotypes of high-throughput CRISPR-Cas9 screens, aggregated gene level CRISPR scores were calculated for each experiments. First, all negative selection screens for cell viability were downloaded from the GenomeCRISPR database (Rauscher et al. 2017). First, all genes targeted by less than 3 sgRNAs and all sgRNA where < 30 counts were observed in the time point 0 (T0) sample, were removed from each screen individually. In addition, we excluded all sgRNAs in the GeCKOv2 library (Sanjana et al. 2014) that were flagged as ‘isUsed = FALSE’ in the ‘Achilles_v3.3.8.reagent.table.txt’ (https://portals.broadinstitute.org/achilles/datasets/7/download) on the Project Achilles (Aguirre et al. 2016) website. After filtering, raw read counts were corrected for differences in sequencing depth by dividing the each read count by the median of all read counts of samples at both T0 and the final time point. Based on these values, fold changes were calculated for technical replicates, after adding 1 to each count to avoid logs of 0, as

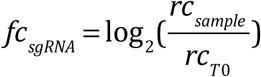

where *rc*_*sample*_ is the normalized read-count measured in the sample cell population and *rc*_*T0*_ is the normalized read count measured at time point 0. In some cases, the read-count abundance in the plasmid DNA pool was given instead of time point 0 sequencing data of cells. In these cases, the plasmid DNA read-counts were used to calculate the fold changes for all sample replicates of those screens. Furthermore, in 2 cases (Doench et al. 2016; Munoz et al. 2016) no read count data was available. Here we used the original fold change values provided by the authors of the experiments.

In order to assess the quality of each screen, Bayesian Analysis of Gene Essentiality (BAGEL; Hart & Moffat 2016) was used to predict gene essentiality. Using precision-recall-curves the ability to separate core-essential and non-essential genes based on the fold change data was examined. All screens where an area under the precision-recall-curve of less than 0.85 was observed were excluded from further analysis. After selecting screens for downstream analysis (Supplementary Table 2), gene level CRISPR scores were calculated as the average fold change of all sgRNAs targeting a gene. We then used quantile normalization to normalize CRISPR scores across experiments.

### Gene level correction of library batch effect

In order to estimate batch effects introduced by the use of different libraries, a robust linear model of the form *y*_*i*_ = β_0_ + β_1_*x*_i1_ +…+ β_n_*x*_in_ + ε_i_ β_0_ = 0 and *y*_*i*_ = *y*_*CRISPR,i*_ – *Median*(*y*_*CRISPR*_) with was fitted for each gene individually where *n* is the number of libraries including the gene, *i* is the index of a data point and *y*_*CRISPR*_ are quantile-normalized CRISPR scores. The coefficients β_1_…β_n_ are then the estimated difference between the CRISPR scores screened in a library to the median CRISPR scores across all libraries. A robust F-test as implemented in the R package ‘sfsmisc’ (Maechler 2008) was used to test the null hypothesis that the median CRISPR score observed for a gene is the same across all libraries. The Benjamini-Hochberg method (Benjamini & Hochberg 1995) was used to estimate the false discovery rate (FDR) for each test. In case the null hypothesis could be rejected at 5% FDR a library specific batch effect was assumed and CRISPR scores observed using that library were centered by subtracting its distance to the median of CRISPR scores across all libraries. A library was flagged from batch correction in cases where a similar (same sign of the model coefficients) batch effect was predicted for the libraries used in the screens of Wang et al. (Wang et al. 2017) and Tzelepis et al. (Tzelepis et al. 2016). Both of these libraries were used to screen primarily acute myeloid leukemia (AML) cell lines and thus the null hypothesis described above might not hold true in the case of AML specific genes. Therefore, in such cases, no batch adjustment was performed.

### Quality control of normalized CRISPR scores

To assess the appropriateness of the normalization steps described above quality control was performed examining several different properties of the normalized data. First of all, samples were clustered to evaluate if biologically related samples clustered more closely than more biologically distant samples. Here, the set of genes shared across all libraries was determined and Ward clustering (as implemented in R’s ‘ward.D2’ method for hierarchical clustering) was performed. The ‘pheatmap’ R package was used to visualize the heat map shown in Figure 2A. Next, differences in normalized CRISPR scores across samples were observed at the examples of 9 core-essential polymerases, and 9 genes situated on the Y chromosome, all of which were sampled randomly from the set of core-essential polymerase genes (Hart et al. 2017) and the set of Y chromosome genes, respectively. Only screens in female cell lines were plotted in Figure 2C. To examine if normalized CRISPR scores could distinguish core-essential genes (Hart et al. 2017) from non-essential genes (Hart et al. 2015) precision-recall-curves were generated for each screen using the ROCR R/Bioconductor package (Sing et al. 2005; Gentleman et al. 2004). Further, a number of control oncogenes (KRAS, NRAS, BRAF and PIK3CA) were selected to see if an expected difference in response to gene knockout depending on the mutation status of the gene could be observed. P-values shown in Figures 1E-H were calculated using a two-sided Student’s t-test as implemented in R. Finally we checked that potential unwanted effects introduced by the batch correction did not distort findings published in the papers where data was included in our pipeline. For these comparisons normalized CRISPR scores were used for the cell lines featured in the original publications.

### Combinatorial testing of gene-gene interactions

To test for differences in fitness response based on loss-of-function genotypes, fitness scores for all CRISPR-Cas9 screens in cell lines were genotype information was available were selected. We selected all genes that were marked as altered by somatic mutations or copy number changes in at least 3 or marked as overexpressed in at least 5 distinct cell lines as query genes. In total, 221 genes were selected. Consequently, we identified all combinations between these query genes and genes perturbed in screens (target genes). Target genes were selected such that, fitness scores were available for at least 3 distinct cell lines with and without a query loss-of-function. Overall, we identified ~3.8 million such combinations. As input data for the test, we used normalized CRISPR scores as described above. We fitted a linear mixed-effects model for each combination, modeling the loss-of-function genotype as fixed effect and the cell line as random effect to account for cell-line-specific biases. For modeling, the R package ‘lme4’ (Bates et al. 2014) was used. The R package ‘lmerTest’ (Kuznetsova et al. 2016) was used to calculate an estimation of significance (p-value) for each model. After testing similar queries were identified by calculating the Pearson correlation of the estimated model coefficients for each pair of query genes. Pairs of query genes with a 100% correlation were merged together into a ‘meta’ query gene. To control the expected fraction of false discoveries made during multiple testing independent hypothesis testing (IHW; Ignatiadis et al. 2016) was used using the variance of the normalized CRIPSR scores of the altered (mutated or overexpressed) group as a covariate for hypothesis weighting (Figure S2C, Figure S2D).

### Quantification of genetic interactions

Interactions between genes were quantified using the π-score statistic (Horn et al. 2011; Laufer et al. 2013; Fischer et al. 2015). π-scores were calculated using the ‘HD2013SGImaineffects’ function implemented in the R/Bioconductor package ‘HD2013SGI’ (Laufer et al. 2013). To generate the input for the ‘HD2013SGImaineffects’ function, normalized CRISPR scores were entered by subtracting column means and scaled by dividing columns by their standard deviation.

### Gene set enrichment network

To generate the gene set enrichment network shown in Figure 3F we selected 10 query genes and all target genes interacting with these queries at FDR < 20%. The resulting list of edges was visualized in Cytoscape (Shannon et al. 2003) using a force-directed spring-embedded network algorithm. Query gene nodes were arranged manually. ConsensusPathDB (Kamburov et al. 2013) was used to perform gene set over-representation analysis and for each query gene a pathway term was selected from the list of results. The q-values displayed in Figure 3F are as provided by ConsensusPathDB. We would like to mention that Figure 3F was inspired by a previous study by M. Brockmann and colleagues (Brockmann et al. 2017).

### TCF4/Wnt-luciferase reporter assay

HEK293T cells were cultured in Dulbecco's MEM (GIBCO) supplemented with 10 % fetal bovine serum (Biochrom GmbH, Berlin, Germany) without antibiotics. Experiments were performed in a 384-well format using white, flat-bottom polystyrene plates (Greiner, Mannheim, Germany). HEK293T cells were reverse transfected with 20 nM indicated siRNAs with the help of 1% of Lipofectamine RNAiMAX Transfection Reagent (#<u>13778150</u>; Thermo Fisher Scientific Waltham, MA, USA). 24 hrs later cells were transfected with 0.2% of TransIT-LT1 transfection reagent (731-0029; Mirus/VWR, Madison, USA) and 20 ng of TCF4/Wnt firefly luciferase reporter, 10 ng of actin-*Renilla* luciferase reporter, and the canonical Wnt signalling was induced by addition of the Wnt3(20 ng)-, β-catenin(20 ng)- or Dvl3(5 ng)- expressing plasmids or left without induction by addition of the Ctrl plasmid pcDNA3. Luminescence was measured with the Mithras LB940 plate reader (Berthold Technologies, Bad Wildbad, Germany). TCF4/Wnt-luciferase signal was normalized to the actin-*Renilla* luciferase reporter signal.

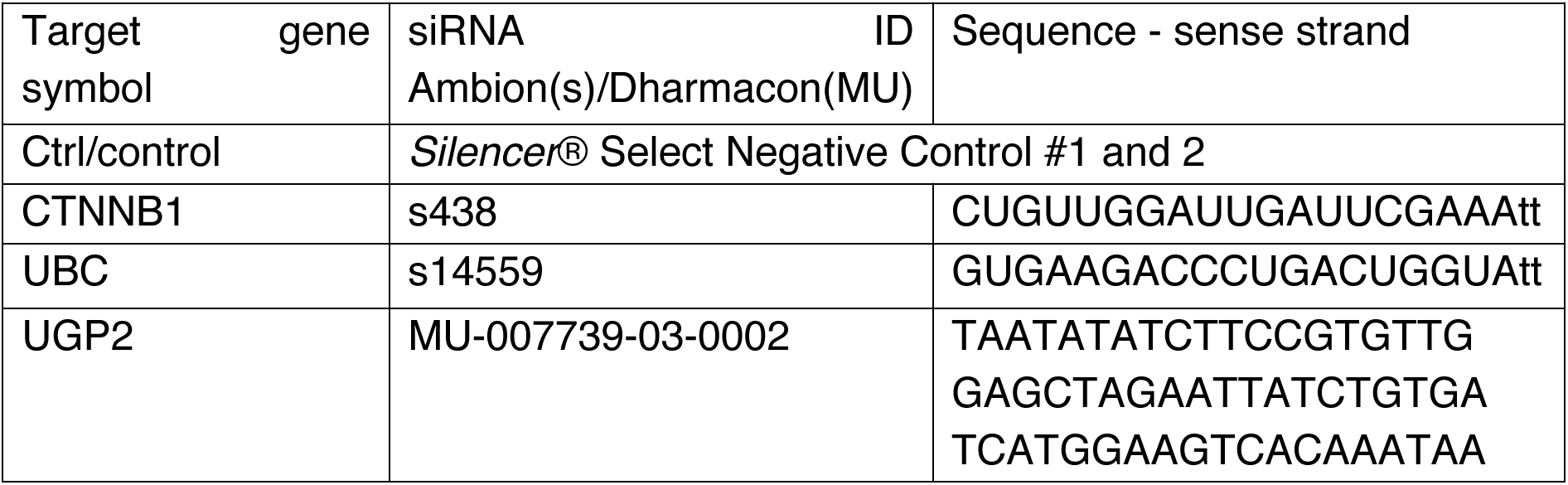
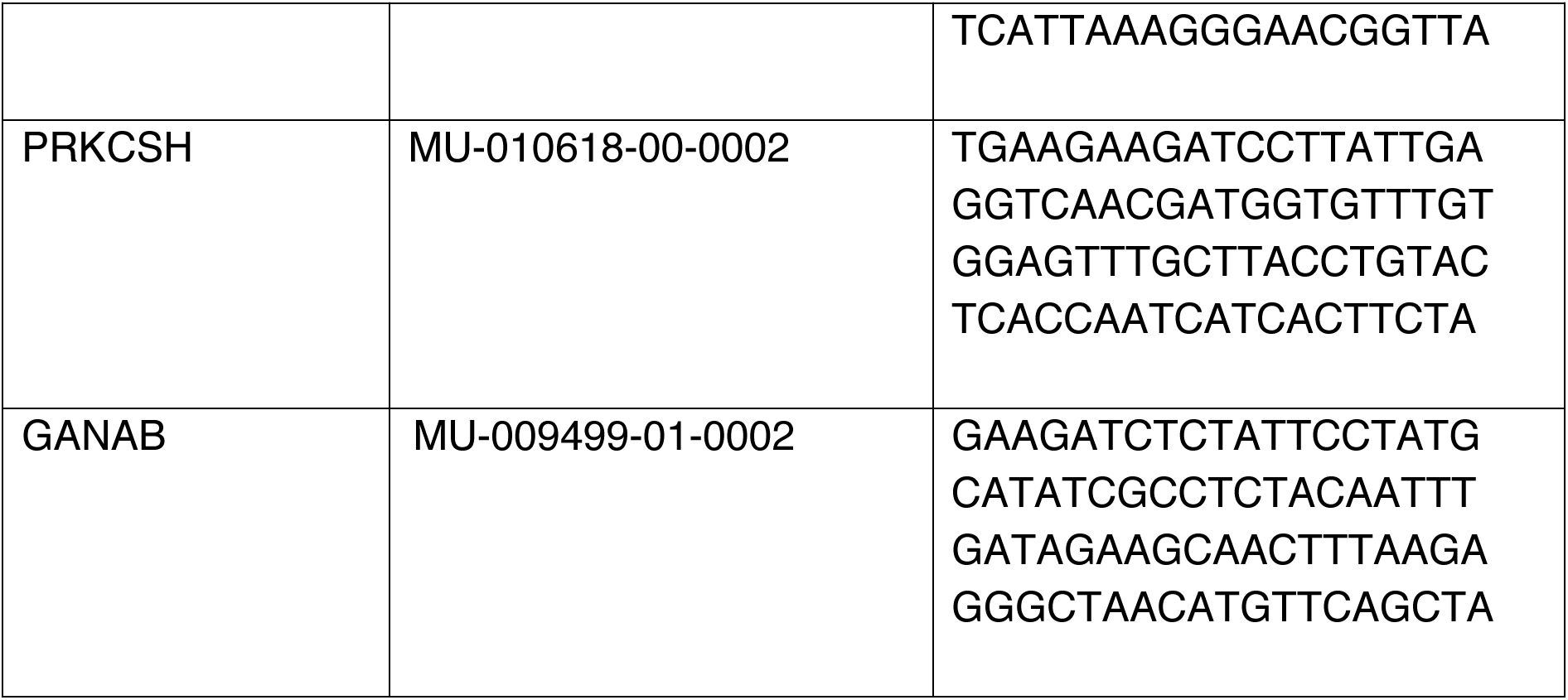
siRNAs

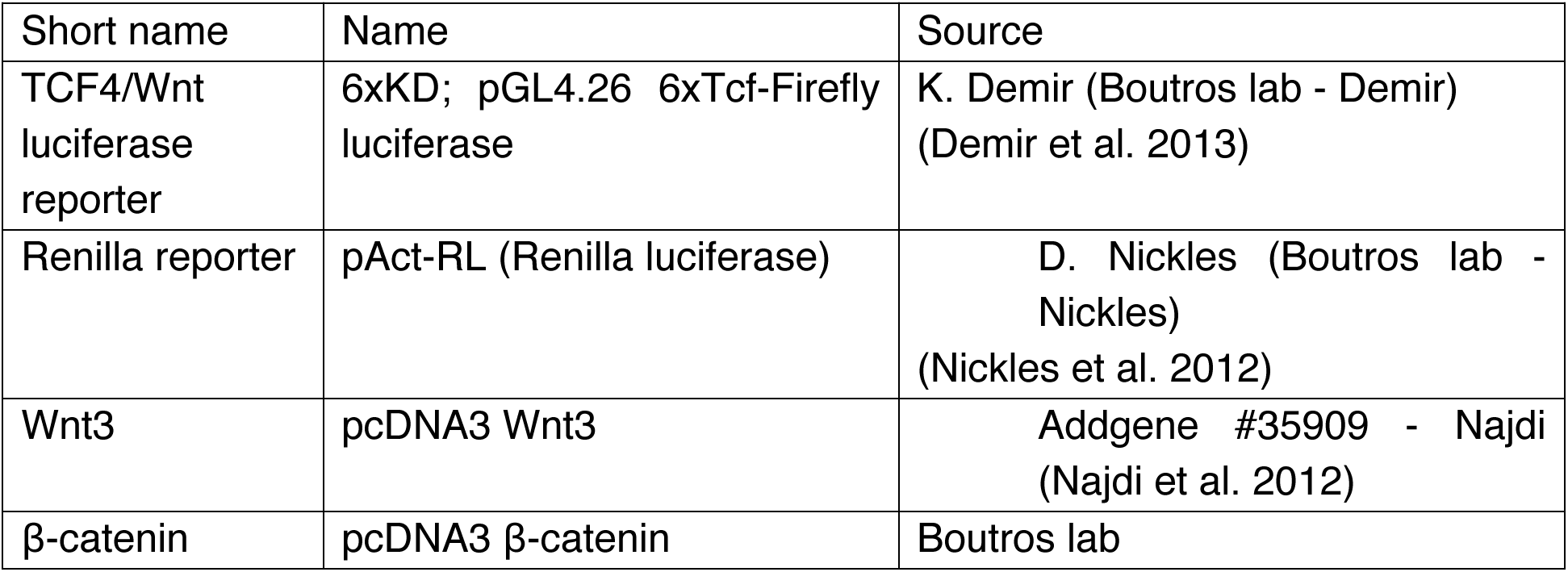
Constructs

Constructs
| Short name | Name | Source |
| --- | --- | --- |
| TCF4/Wnt luciferase reporter | 6xKD; pGL4.26 6xTcf-Firefly luciferase | K. Demir (Boutros lab - Demir) (Demir et al. 2013) |
| Renilla reporter | pAct-RL (Renilla luciferase) | D. Nickles (Boutros lab - Nickles) (Nickles et al. 2012) |
| Wnt3 | pcDNA3 Wnt3 | Addgene #35909 - Najdi (Najdi et al. 2012) |
| β-catenin | pcDNA3 β-catenin | Boutros lab |

### NanoLuciferase Wnt3 secretion assay

Similar to the TCF4/Wnt-luciferase reporter assay, HEK293T cells were reverse transfected with indicated siRNAs and seeded into 384-well format white, flat-bottom polystyrene plates (Greiner, Mannheim, Germany). 24 hrs later cells were transfected with 20 ng of NLucWnt3 or Wnt3NLuc expression constructs (NanoLuciferase sequence; Hall et al. 2012) was cloned into pcDNA Wnt3 plasmid either after the signal peptide or at the C-terminus) together with 5 ng of CMV Firefly luciferase reporter plasmid. 48 hrs later the plates were centrifuged and 20 μl of culture medium was transferred to a new plate. NanoLuciferase signal in the lysate and medium was detected with the help of a Nano-Glo Luciferase Assay (#N1110) from Promega (USA) according to the manufactures instructions. Luminescence was measured with the Mithras LB940 plate reader (Berthold Technologies, Bad Wildbad, Germany). In the case of the lysate, first the signal for Firefly luciferase and then for NanoLuciferase was measured. The NanoLuciferase signal in the culture medium was normalized to the NanoLuciferase signal in lysate normalized to the Firefly luciferase signal.

Constructs

Constructs
| Short name | Name | Source |
| --- | --- | --- |
| TCF4/Wnt luciferase reporter | 6xKD; pGL4.26 6xTcf-Firefly luciferase | K. Demir (Boutros lab - Demir) (Demir et al. 2013) |
| Renilla reporter | pAct-RL (Renilla luciferase) | D. Nickles (Boutros lab - Nickles) (Nickles et al. 2012) |
| Wnt3 | pcDNA3 Wnt3 | Addgene #35909 - Najdi (Najdi et al. 2012) |
| β-catenin | pcDNA3 β-catenin | Boutros lab |
| NLucWnt3 | pcDNA_NLucWnt3 | Boutros lab |
| Wnt3NLuc | pcDNA_Wnt3NLuc | Boutros lab |
| CMV firefly luciferase | pLenti CMV Firefly luciferase | Campeau et al. 2009 |

### Gene similarity network benchmarking and modeling

In order to assess whether interaction similarity networks can predict protein complex co membership, protein complex annotations were downloaded from the CORUM databases (Ruepp et al. 2010) and target genes included in the CORUM data were selected. We removed all pairwise interactions π_*tq*_ with |π_*tq*_| < π_min_ where π_*tq*_ is the interaction score between target gene *q* and π_min_ query gene *q* and min is a chosen threshold. Subsequently, the Spearman correlation was calculated as implemented in the ‘Hmisc’ R package for each possible pair of target genes using pairwise complete observations. Target gene pairs where less than *n*_min_ data points were used to calculate the correlation were excluded. This analysis was performed for 6 different combinations of the parameters π_min_ and *n*_min_ and ROC curves were drawn to visualize how well the resulting correlations could predict protein complex co-membership as annotated in CORUM. Based on these results π_min_ = 0.2 and *n*_min_ = 15 were selected as thresholds to calculate Spearman correlations between all possible target gene pairs as described above. For each correlation the asymptotic p-value was computed using the ‘Hmisc’ R package. Bonferroni correction was applied to the resulting p-values and gene pairs with an adjusted p-value < 0.5 were used as edges for the gene similarity network in Figure 5C. The network was visualized using Cytoscape (Shannon et al. 2003). A force-directed spring-embedded layout was used to position the nodes of the network without edge weighting. The visual representation of the network was inspired by previous studies in yeast (Costanzo et al. 2016; Costanzo et al. 2010). The Spatial Analysis of Functional Enrichment (SAFE; Baryshnikova 2016b; Baryshnikova 2016a) Cytoscape-plugin was used to identify functional modules in the network. For SAFE analysis, the map-based distance-metric was chosen with a maximum distance threshold of 0.6 (percentile). To build the composite map, a minimal landscape size of 7 was chosen and the Jaccard distance was used as a similarity metric for group attributes with a similarity threshold of 0.75. As background for the enrichment, all nodes in the annotation standard were chosen. In SAFE the annotation standard is a binary matrix of genes (rows) and annotation terms (columns). A value of 1 indicates that a gene is annotated with a specific annotation term. For our analysis, we generated such an annotation standard containing Gene Ontology (GO; Ashburner et al. 2000) Biological Process annotations for all target genes tested. GO annotations were downloaded from the example data section of the SAFE algorithm’s GitHub page (https://github.com/baryshnikova-lab/safe-data/blob/master/attributes/go_Hs_P_160509.txt.gz; accessed 13/09/2017) and filtered to contain only genes tested in our interaction analysis.

## Acknowledgements

We thank Niklas Rindtorff, Tianzuo Zhan, Johannes Betge and Christian Scheeder for critical comments on the manuscript. We would like to thank the Boutros lab for helpful discussions.

## Author contributions

BR, FH, and MB designed the study. BR wrote the analysis code. TH consulted on statistical analysis. LH and OV performed the experiments. All authors discussed and analyzed results. BR, FH, LH, OV and MB wrote the manuscript. All authors read and approved the final manuscript.

## Conflict of interest

We declare no conflict of interest.

## Funding

B.R. was supported by the BMBF-funded Heidelberg Center for Human Bioinformatics (HD-HuB) within the German Network for Bioinformatics Infrastructure (de.NBI) (Grant #031A537A). Work in the Boutros lab is supported in part by an ERC Advanced grant.

